# Research challenges in Africa – an exploratory study on the experiences and opinions of African researchers

**DOI:** 10.1101/446328

**Authors:** Lem Ngongalah, Wepngong Emerson, Ngwa Niba Rawlings, James Muleme Musisi

## Abstract

**Background:** Insufficient research is a major impediment to growth, development and advancement of health in Africa. Africa produces less than 1% of global research output. Meanwhile, African countries face some of the toughest challenges worldwide, most of which can only be tackled through robust and efficient research. Addressing the barriers to conducting research in Africa is a step towards improving research capacity and output. This study aimed to identify the key challenges affecting research practice and output in Africa; and to highlight priority areas for improvement.

**Methods:** A cross-sectional survey was administered through an online questionnaire, including participants from six countries in Sub-Saharan Africa. Participants included research professionals, research students, research groups and academics.

**Results:** A total of 424 participants responded to this survey. The ability to conduct and produce high-quality research was seen to be influenced by multiple factors, most of which were related to the research environment in African countries. Priority areas for improvement included providing more training, raising awareness on the importance of research in Africa, encouraging governments to commit to research and increasing collaboration between researchers in Africa.

**Conclusion:** The conditions under which research is done in Africa are severely flawed and do not encourage engagement in research, or continuity of research activity. African governments need to develop initiatives that accelerate and support research and research-based education in Africa, in order to build a solid foundation for research, increase research capacity, and enable institutions to provide valuable training and develop sustainable research opportunities in Africa.

## 1. Introduction

The African continent is disproportionately plagued by a multitude of diseases, many of which have been eradicated in other parts of the world. Africa faces some of the toughest challenges worldwide, some of which include poor disease management strategies, poor infrastructural development, food insecurity, poor hygiene and sanitation, lack of potable water and climate change hazards [1]. Tackling such challenges requires robust and efficient research. Research is key to the development of any nation, and a significant determinant of health and productivity [2]. Research is also important for the generation of knowledge useful for policy-making, planning and strategic action, all of which are critical to Africa’s development [3,4]. Africa contributes less than 1% of global research output [5]. This is in contrast to the situation in most high income countries where research is receiving increased attention and investment, in recognition of its acknowledged contribution to health and economic development [6]. Insufficient research in Africa translates into data gaps, which are a major constraint to the creation and implementation of health or development programs. This also brings a challenge to the development of efficient therapies for common diseases, especially those specific to the African continent [7, 8]. As a consequence, several health needs of the African population remain unmet. The paucity of accurate and reliable data further affects the effective monitoring and evaluation of existing programs. For example, without systematically measured estimates on health outcomes and other socio-economic factors in African countries, it is difficult to generate evidence on the effectiveness of existing programs and policies [9]. In the absence of vigorous and high quality research, diseases may continue to resurface in Africa, standard of living will depreciate, death rates will multiply and the African continent will remain under-developed.

While conducting and producing research successfully may be attributed to the competency or hard work of individual researchers or research teams, other factors such as the research environment, availability of facilities and supportive structures all have a role to play. Research structures in most African countries are not resilient enough to address the existing challenges, and the conditions under which research is carried out are severely compromised [7,10]. Addressing the barriers to conducting research in Africa is a step towards improving research capacity and output. The findings of this study will be instrumental in identifying opportunities for improvement, as well as elements that need to be factored into the development of programs and interventions aimed at increasing research output and developing research-friendly environments in Africa. This study therefore aims to explore research challenges in Africa, and to highlight priority factors that need to be improved in order to increase research output in Africa.

## 2. Methods

### 2.1. Study methodology and sampling

This study was a cross-sectional survey including participants from Sub-Saharan Africa (SSA). SSA was purposively selected as it has been shown to have the least number of researchers per million inhabitants globally, compared to other regions [11]. A multi-stage sampling strategy was used to identify participants for this survey. The first stage involved a random selection of English-speaking countries in SSA, using a lottery selection method. All countries in SSA were assigned a unique identification number unknown to the research team. Five members of the research team were then asked to randomly select two countries each. The ten countries corresponding to the numbers selected formed the study sample. Each member then identified participants from their selected countries using purposive, convenience and snowball sampling methods.

This study did not require ethical approval because it was a non-invasive study which did not involve any sensitive information, aspects of vulnerability or risk of harm to participants. The study was completely anonymised and no personal identifiable information was collected. Information on the study protocol, aims, objectives and how participant responses will be used was circulated to all participants. Informed written consent was provided prior to participation.

### 2.2. Questionnaire design

An online researcher-developed questionnaire was used consisting of 10 questions reflecting the contents of a consultative workshop conducted by the Collaboration for Research Excellence in Africa (CORE Africa), on the subject of research challenges in Africa. Details of the workshop are described elsewhere [12]. Briefly, experienced researchers from various African countries were engaged in a series of interviews and group discussions on the state of research in Africa, and the factors that influence research practice and output. A report was generated from this workshop, and the findings served as the basis for the survey questionnaire; in addition to information obtained from previous literature [13-16]. The questionnaire was developed using a Google form.

The questionnaire was piloted prior to the survey, with researchers from two African countries and two members of CORE Africa, who were not included in the study. Piloting was done to assess for accessibility, question wording, participant understanding and other practical issues. The questionnaire was then modified accordingly. A link to the questionnaire was sent to all identified participants by email and through social media platforms such as LinkedIn, WhatsApp and Facebook. Participants included research professionals, research students, research groups and academics. Responses were anonymized and questionnaires were set to accept only one response from each participant.

The questionnaire collected data on the following areas: participants’ countries of origin, research experience, whether and how research training was acquired, motivation for doing research, research challenges in Africa, barriers to conducting research in Africa, factors affecting research output in Africa, awareness of research training institutions in Africa and opportunities for improvement. The survey took approximately 5 – 10 minutes to complete.

### 2.3. Data analysis

Participants’ responses were pooled and analyzed using Microsoft Excel and the Statistical Package for Social Sciences (IBM SPSS Statistics). Data analysis mainly involved descriptive statistics, including frequencies, graphs and tables. The survey responses were further coded and organised into key themes informed by the study objectives and survey questions. Coding and thematic organisation was done by the lead researcher, and then verified by two co-authors for rigour and reliability.

## 3. Results

### 3.1. Participant characteristics

#### 3.1.1. Participant countries of origin

The ten countries selected to participate in this survey included Cameroon, Nigeria, Ghana, Uganda, Tanzania, Kenya, Malawi, Democratic Republic of Congo, South Africa and Rwanda. Participant response rate could not be determined due to the methods through which participants were identified. Participation rate is therefore described based on the countries that were selected to participate. Six (60%) out of the ten selected countries participated in the survey; yielding a total of 424 participants (table 1). No participants responded in four (40%) countries. A majority of responses were from Cameroon (45%) and Nigeria (32%), while the proportion of participants from other countries ranged between 3 – 11%.

**Table 1:** Countries of origin and proportion of survey respondents

| Selected to participate | Participated | n(%) of participants<br>N=424 |
| --- | --- | --- |
| Cameroon | Yes | 191 (45.0) |
| Nigeria | Yes | 136 (32.0) |
| Kenya | Yes | 46 (11.0) |
| Uganda | Yes | 21 (5.0) |
| Tanzania | Yes | 17 (4.0) |
| South Africa | Yes | 13 (3.0) |
| Democratic Republic of Congo | No | 0.0 |
| Ghana | No | 0.0 |
| Malawi | No | 0.0 |
| Rwanda | No | 0.0 |

#### 3.1.2. Participants’ research experience

##### a) Paid research experience

408 participants provided data on paid research experience. About half (49.1%) of the participants had been in paid research positions (Fig 1a). Of these, 30.4% had less than 1 year of paid research experience, 12.8% had up to 5 years of paid research experience and 5.9% had over 5 years of paid research experience. Half of the participants (50.9%) had no paid research experience.

**Fig 1a:**
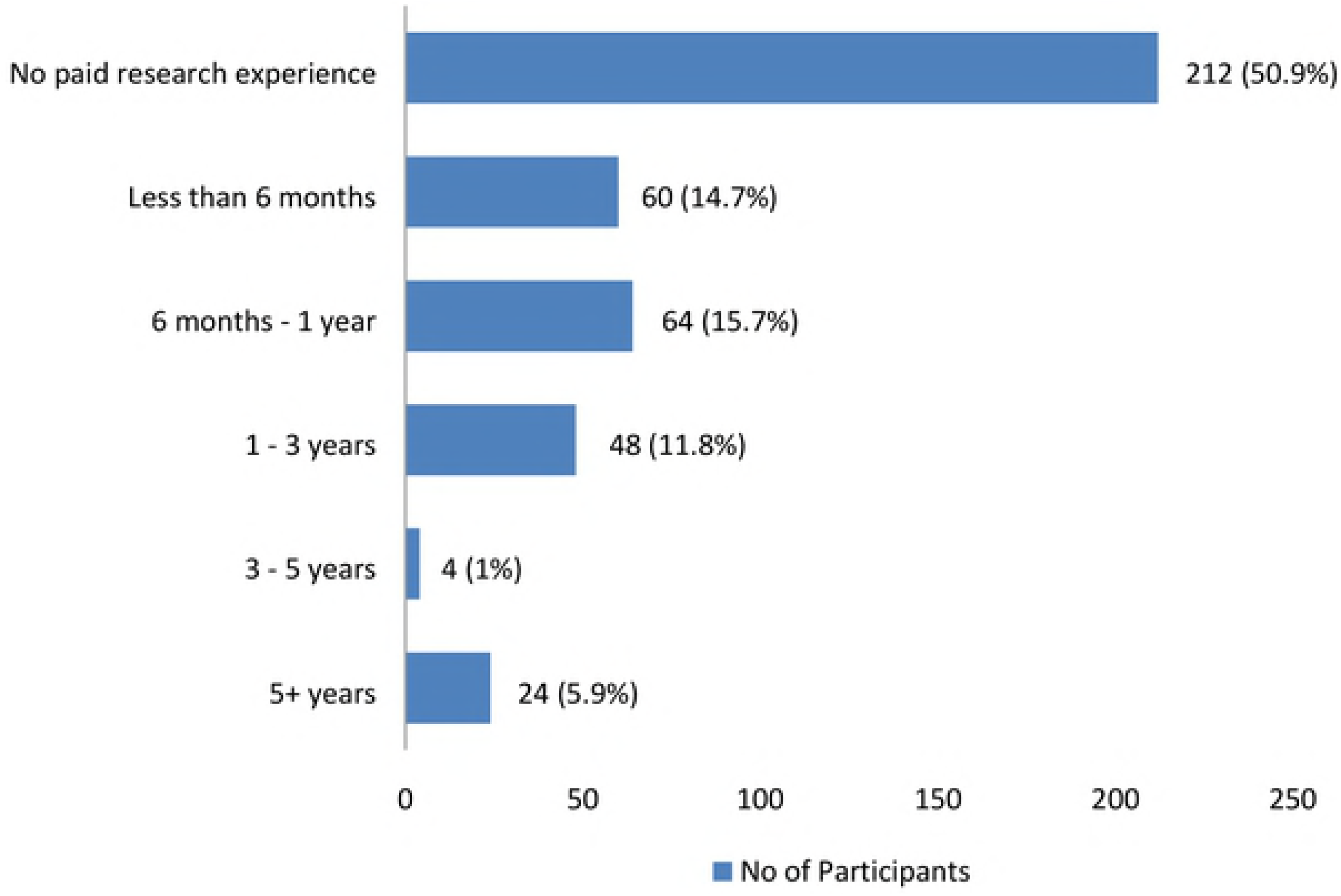
Paid research experience.

##### b) Unpaid research experience

412 participants provided responses on their unpaid (voluntary) research experience. A majority (84.5%) had been in research positions that were unpaid (Fig 1b); 47.6% had less than 1 year of unpaid research experience, 33% had up to 5 years of unpaid research experience and 3.9% had over 5 years of unpaid research experience. 15.5% of participants had no unpaid research experience.

**Fig 1b:**
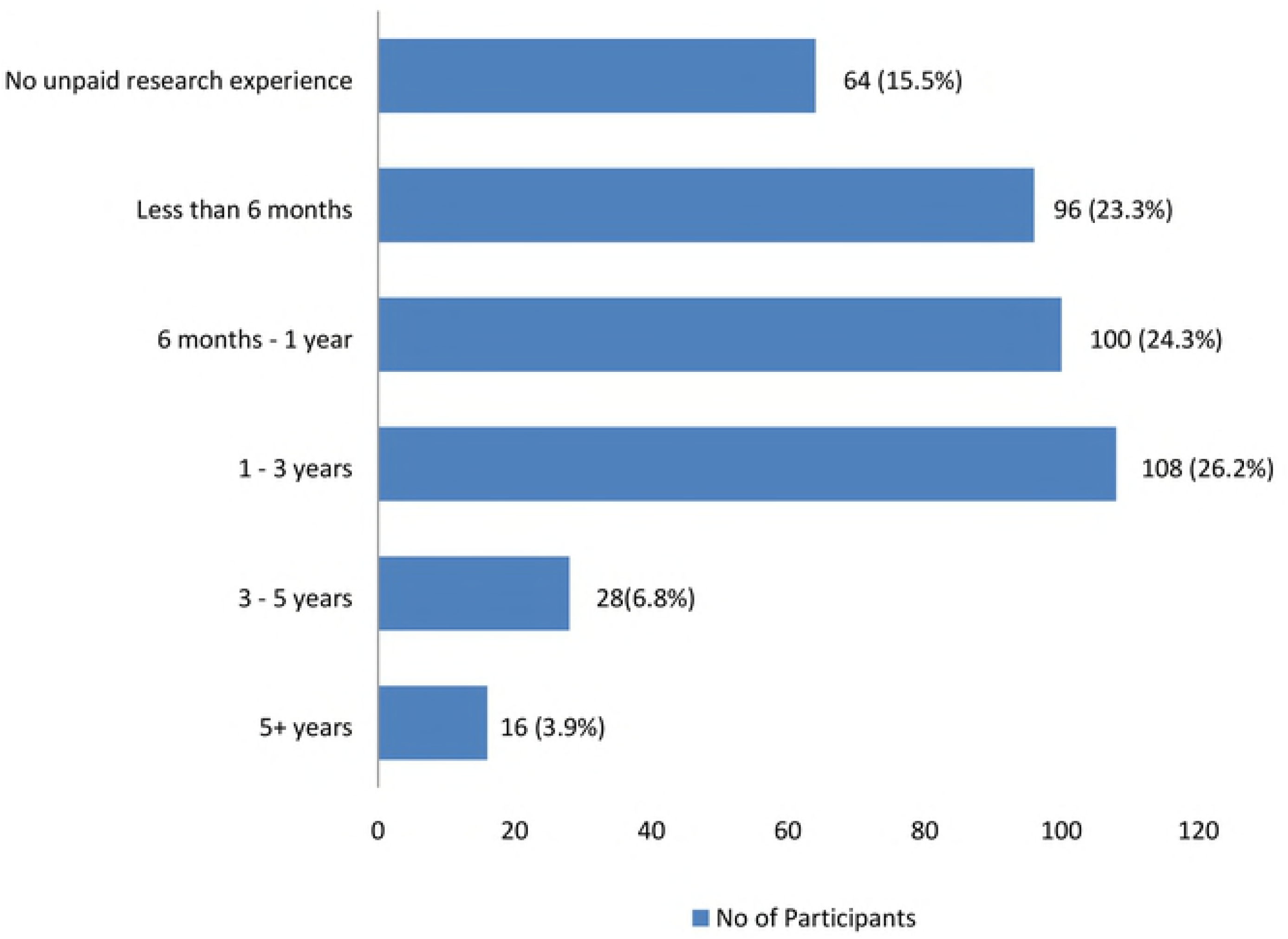
Unpaid research experience.

#### 3.1.3. Sources of research training

416 participants provided responses on where or how they had obtained their training in research. Most participants (76%) developed their research experience through a University or other academic institution (Fig 2). Other sources of research training included internships (24%), paid projects or jobs (14.4%) and research training institutions (6.7%). 10.6% of the participants indicated that they had never received any formal research training.

**Fig 2:**
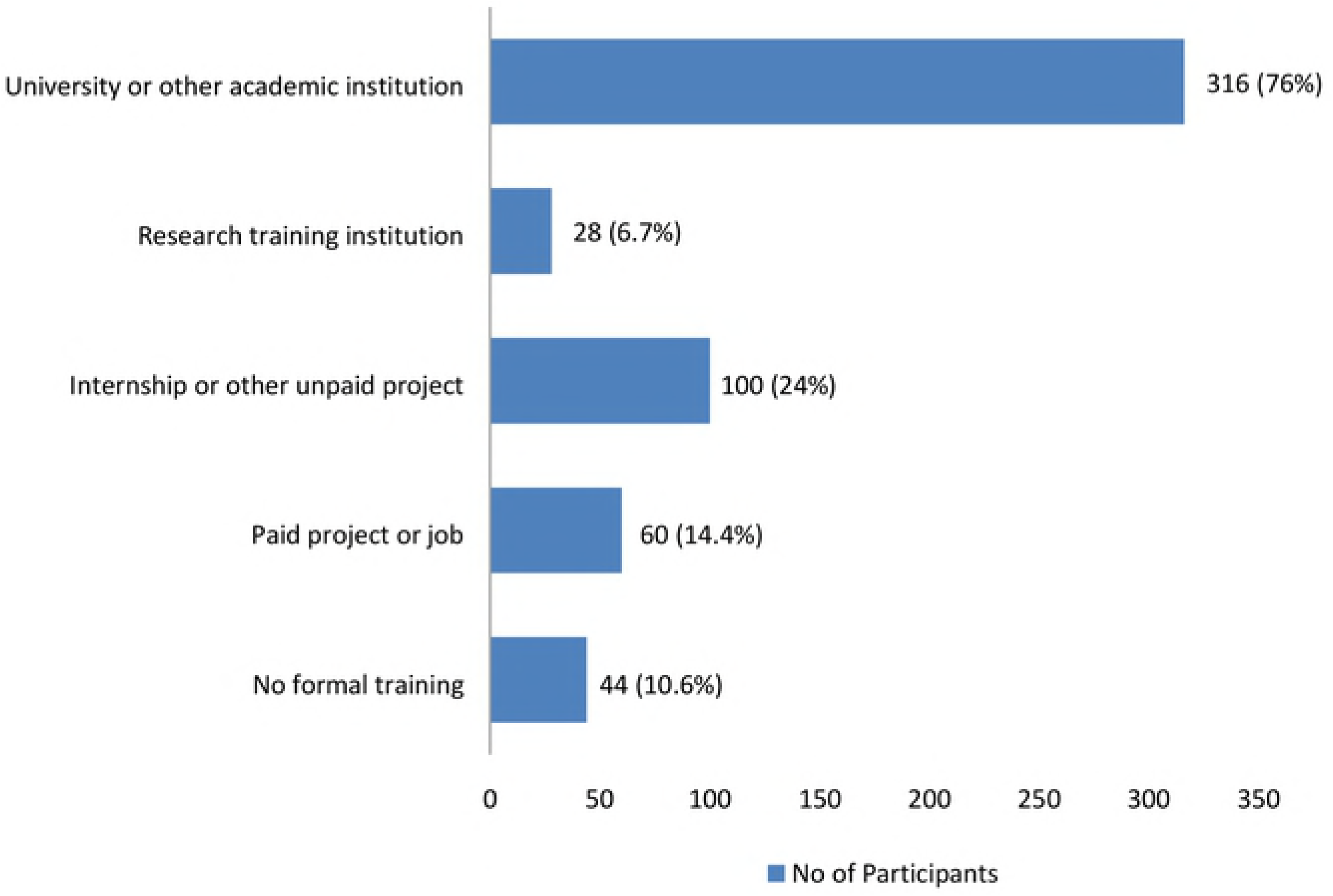
Sources of research training.

#### 3.1.4. Motivation for doing research

420 participants stated the reasons why they did research. The main factor that led participants into the research field was their career choice (68.6%) (Fig 3). This was followed by personal interest in research (37.1%), funded research opportunities (12.4%), being unemployed and wanting something to do (5.7%), and doing a research thesis as a requirement for attaining a University qualification (1%).

**Fig 3:**
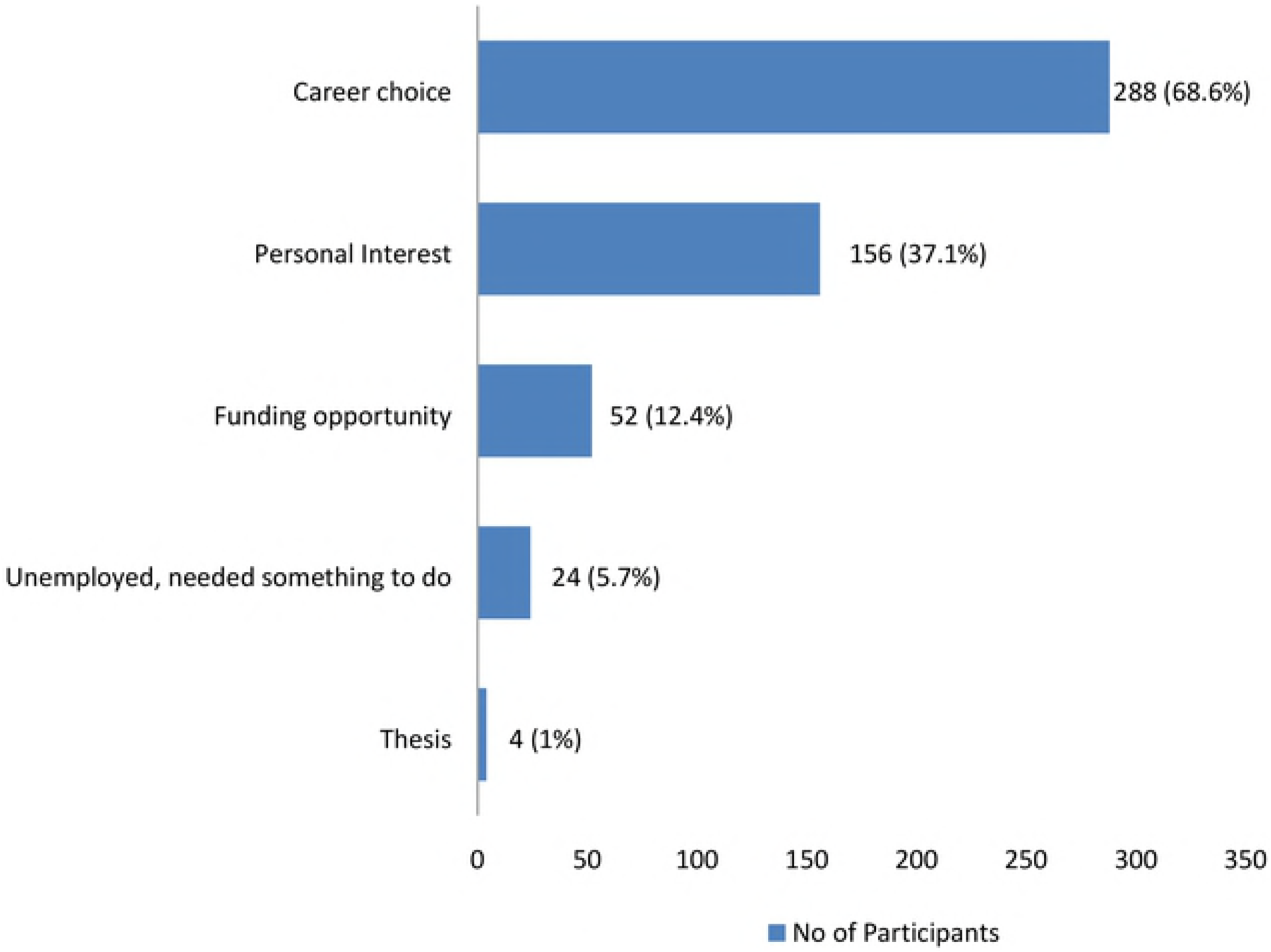
Factors that led participants into doing research.

### 3.2. Research challenges

Three questions explored the participants’ opinions on key research challenges in Africa, the barriers to conducting research and the factors that influence research output in Africa. 420 participants responded to these questions. Participants were asked to select their top three responses for the first two questions; and the top five responses for the third question.

#### 3.2.1. Key challenges affecting research in Africa

The three key research challenges highlighted in this survey included; lack of resources to carry out research, lack of interest or motivation to do research and low uptake of research by African governments and policy makers (Fig 4a). Other factors listed by participants included poor research supervision and lack of motivated research supervisors.

**Fig 4a.**
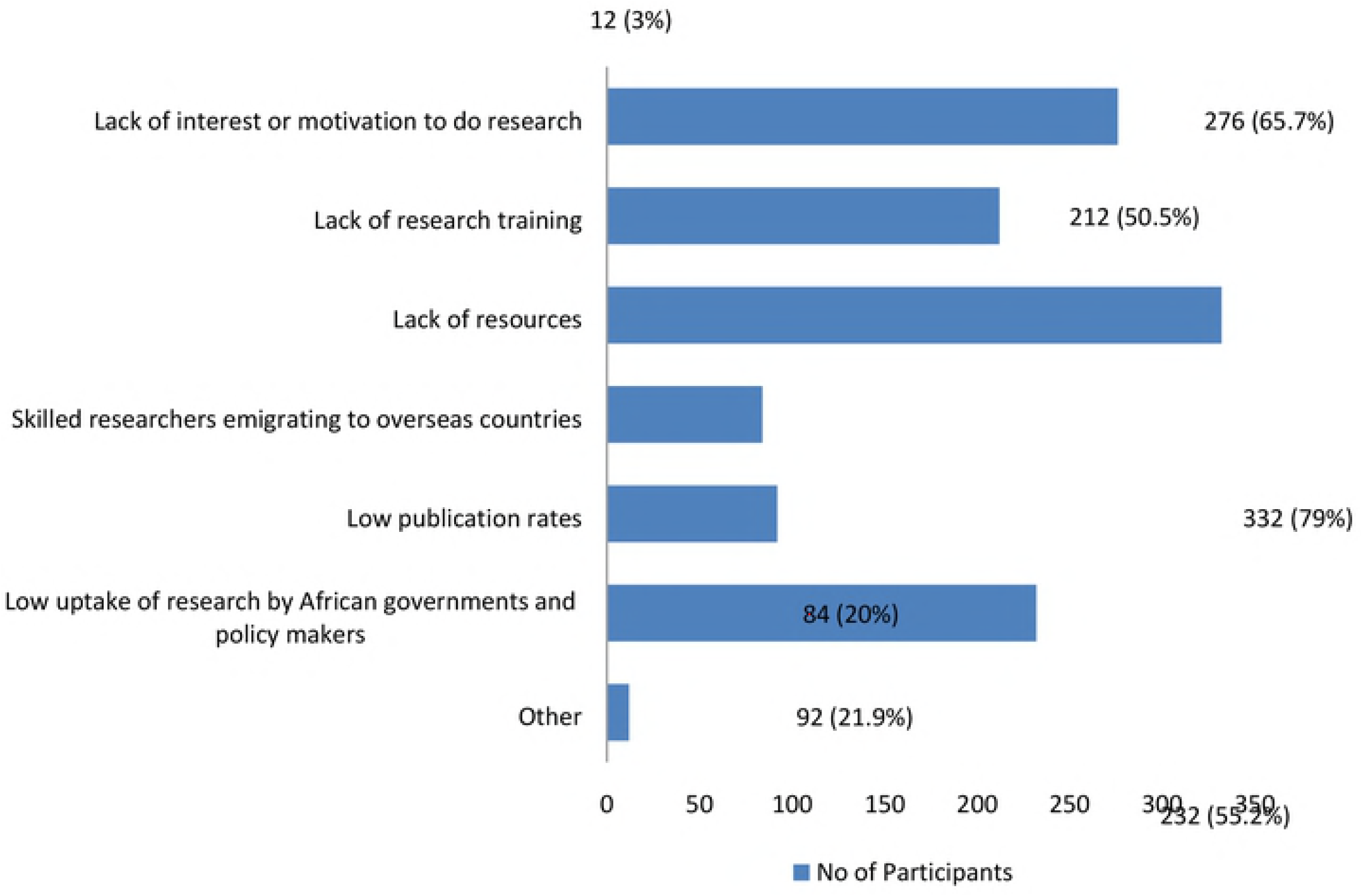
Key research challenges in Africa.

#### 3.2.2. Barriers to conducting research in Africa

The three main barriers to conducting research in Africa were; lack of funding, lack of training facilities and loss of interest or motivation to continue research (Fig 4b).

**Fig 4b.**
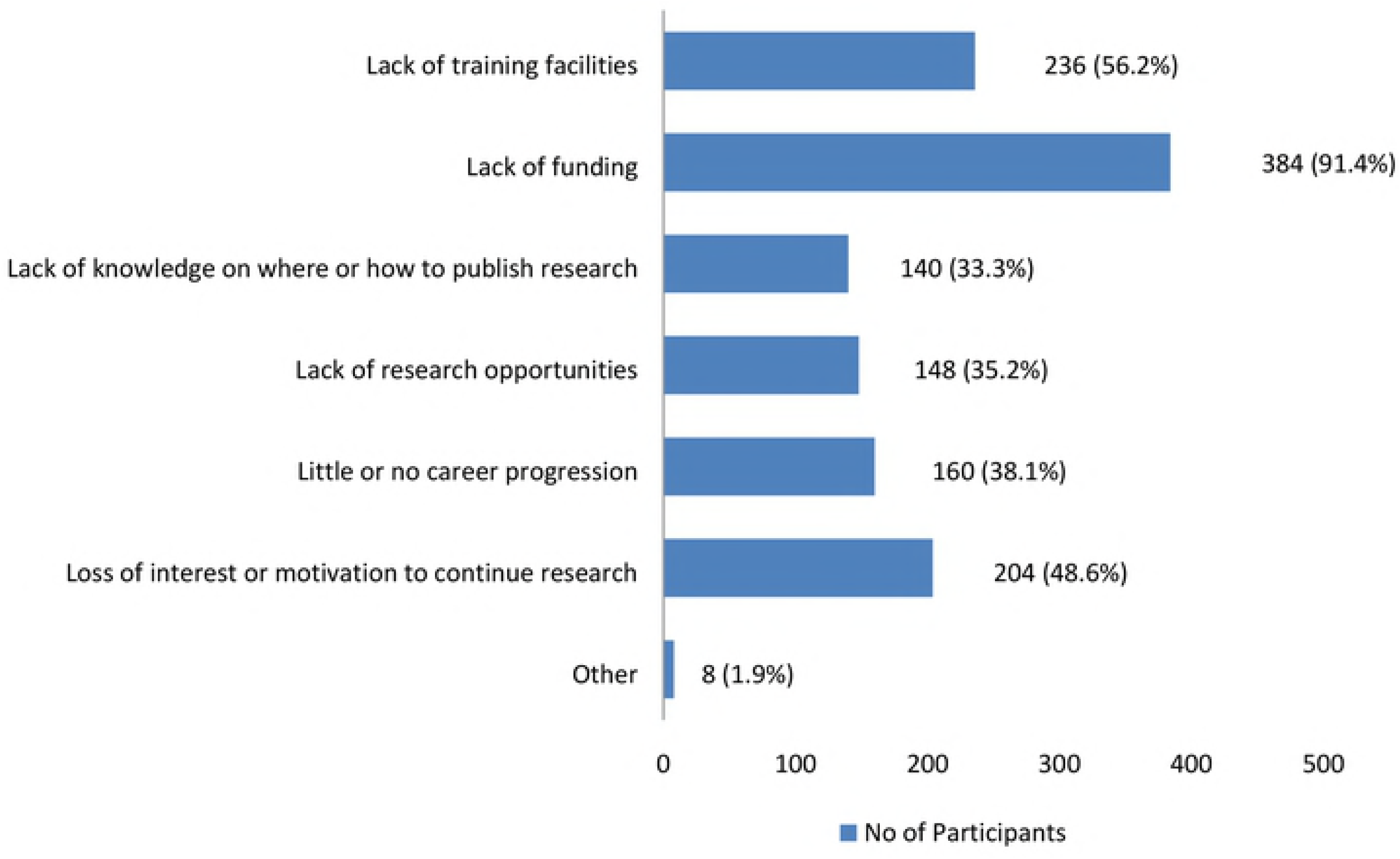
Barriers to the practice of research in Africa.

#### 3.2.3. Factors affecting research output in Africa

The top five responses highlighted as factors which influence research output in Africa, leading to the production of very little scientific research knowledge included; lack of interest or commitment to research by African governments; the lack of facilities to conduct good quality research; the emigration of skilled African researchers from African countries in search for better opportunities overseas; little or no understanding of the importance of research by Africans; and the fact that research priorities for Africa are usually decided by funding organisations from overseas (Fig 4c).

**Fig 4c.**
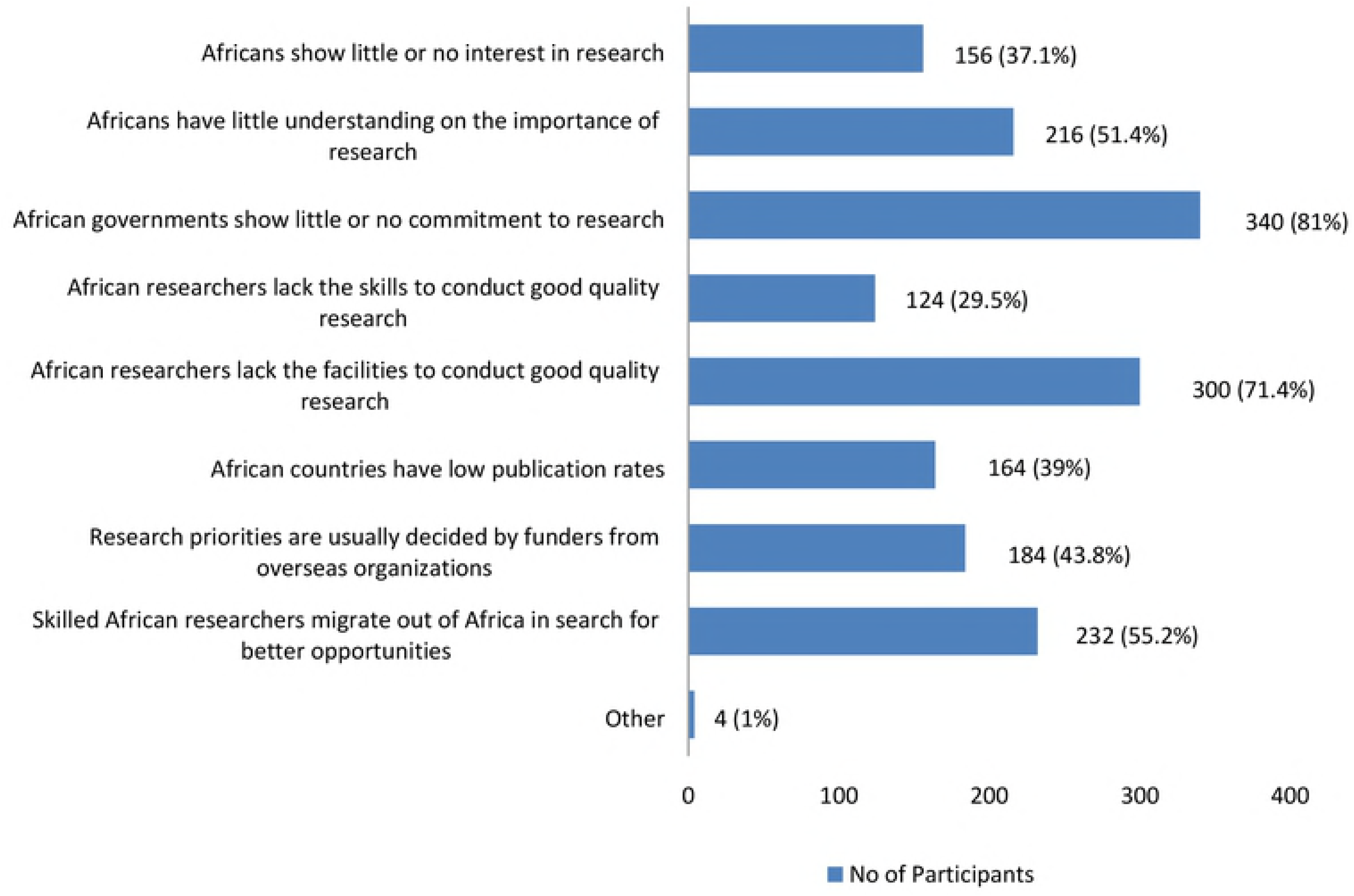
Factors affecting research output in Africa.

#### 3.2.4. Facilities for research training and skill development

Participants were asked to indicate if they were aware of any research institution or organisation providing research training and researcher development programs in their respective countries. A majority of participants (65%) were not aware of any such facility. 35% of the respondents indicated that they were aware of one or more research institutions in their home countries, most of which were in South Africa and Kenya; and a few in Nigeria, Uganda and Cameroon.

### 3.3. Opportunities for improvement

One question explored the participants’ opinions on factors that could help improve research production and output in Africa. Participants were asked to select their top three responses. 420 participants responded to this question. The three most selected responses were: providing training to enable African researchers to produce and publish high quality research; encouraging governments to invest more in research; and raising awareness on the importance of research in Africa (Fig 5). Although not amongst the top three choices, almost half of the survey respondents (44.8%) also indicated that increasing collaborations amongst researchers could help improve research production and output in Africa.

**Fig 5:**
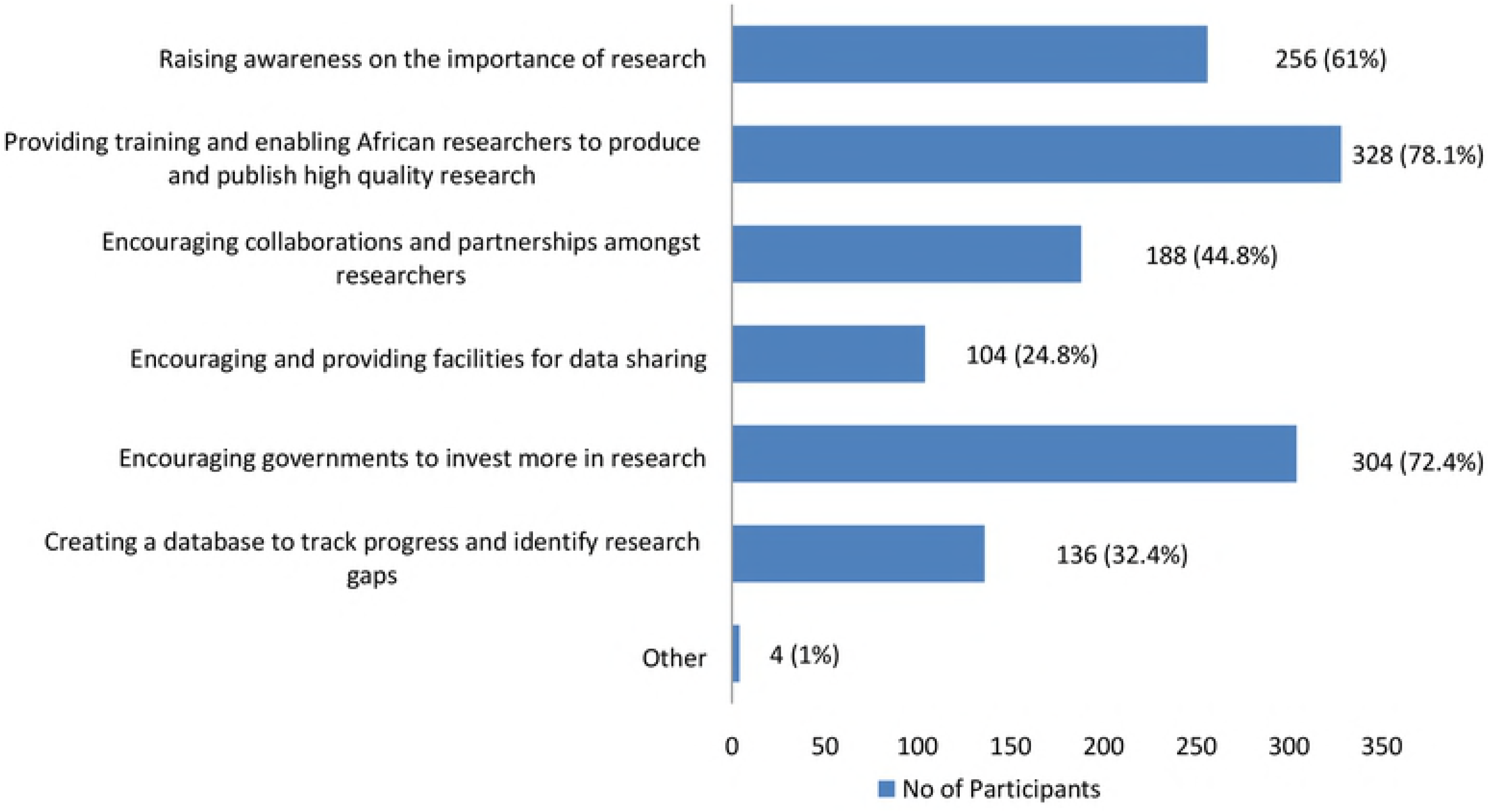
Areas for improvement.

The final part of the survey asked participants to rank three factors on a scale of 1 to 3 in order of importance (highest to lowest), as priority areas that need to be considered in order to improve research output in Africa. These factors included funding, training and support; and collaboration. 420 participants responded to this question. For first priority, the highest number of responses (49%) indicated providing more training and support. In second place, a majority of the respondents (41%) selected more funding. Increasing collaboration was the third priority (Fig 6).

**Fig 6:**
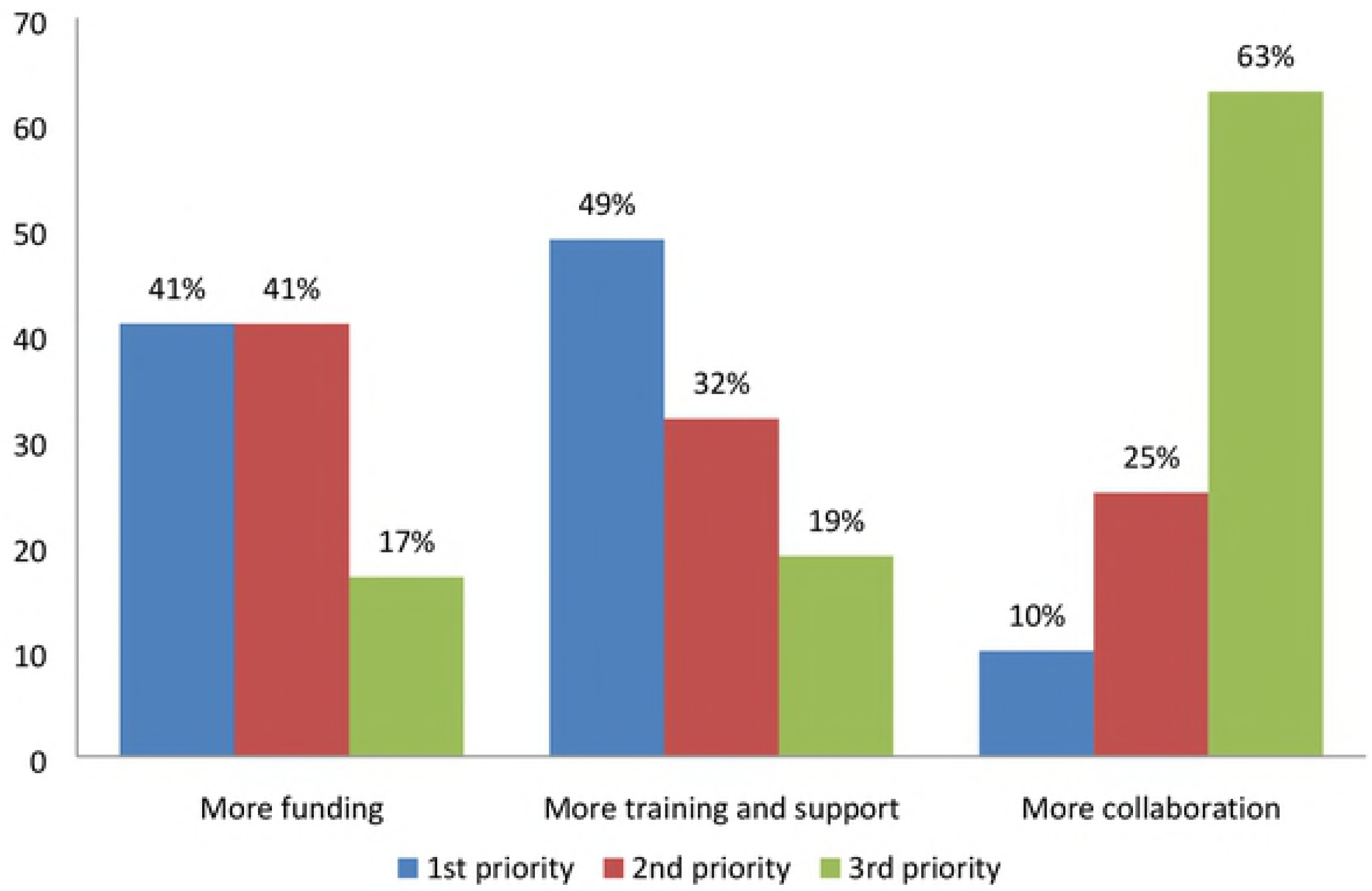
Priority areas for improvement.

## Discussion

This study aimed to identify key research challenges in Africa, as well as prospective areas which if addressed, could potentially increase research production and output in Africa. The emergent challenges from this survey can be clustered into two themes; individual and environmental factors.

Individual factors refer to the challenges that prevent researchers from making the decision to do or continue doing research. The challenges reported at this level included the lack of interest or motivation to do research. These challenges have also been reported in other studies [7,15,17]. The development of interest in research and individual ability are strongly rooted in the quality of education obtained. An educational system that encourages curiosity and promotes the pursuit of further knowledge or ideas fosters the development of a positive research culture [18]. The lack of such a foundation in African countries may partly explain the lack of interest to do research. Sustaining interest in research requires guidance, as well as motivation to continue. Research guidance can be provided through the prior or concurrent work of other researchers in related fields, or through active guidance from experienced researchers [18]. For example, three studies [7,10,15] have found that after developing interest in research, researchers needed further support in the form of mentorship from senior researchers and advice or encouragement from peers. The emigration of skilled researchers out of Africa as shown by this survey, thereby creates a challenge for current and aspiring researchers.

Motivation to continue research on the other hand is determined by factors such as the uptake and use of research findings and ideas, as well as recognition of achievement [19]. The low uptake of research findings by African governments and policy-makers and their lack of commitment to research as reported in this survey do not create a motivating atmosphere; hence, the lack of motivation to do research. One study also found this to be a reason why researchers emigrate to other countries where their efforts are better rewarded [19].

Environmental factors refer to the research environment, which can be described as the structures, legislations and support systems put in place for the use of research. The challenges under this theme are factors relating to research institutions and government bodies. The capacity of individual researchers, including skills, competencies and attitudes to research, are primarily developed through organised training programs and other research activities. This survey found that most participants with research experience obtained their training through Universities and other academic institutions. However, the lack of research training was still a major challenge, and a majority of participants were unaware of any research training institutions in their countries. This is either an indication of a lack or inadequacy of research training programs in African universities. Without institutions with adequate systems for training, guidance and supervision, it is impossible for researchers to develop the skills and expertise required to produce and publish good quality research. It is therefore imperative that African universities become more research-intensive. Although African universities have been marginal in their contribution to global scientific knowledge, universities in some countries are working hard to turn the tide [20]. An overall challenge that remains at this level is the fragile funding structures for research in African universities, which limits the sustainability of research training efforts.

When it comes to research funding, African governments have a great role to play. However, research is often undervalued by government officials and policy-makers as a means to address challenges in Africa. This is evident in their lack of commitment and minimal support for research. Most African countries reportedly contribute very little to research and development [21]. Increasing support for research efforts in Africa will help develop a strong knowledge base, to help inform the development of efficient evidence-based policies and improve growth and development in Africa. African governments can increase their support for research by generating new funding schemes for research training and research-based education in Africa. This will also help increase the capacity of African universities and research institutions to provide adequate training and develop sustainable research opportunities in Africa.

One of the strengths of this report was the inclusion of multiple African countries, to account for international differences in opinions, given that research engagement varies across countries. The study is however limited by the small sample size, the recruitment methods used which are subject to bias and the fact that the questionnaire was only available in English language.

## Conclusion

This study identified key research challenges in Africa, and potential areas for improvement. The emergent themes involved factors at the level of individual researchers, research institutions and government bodies. The research environment in Africa was seen to be the major inhibitor for research in Africa. Research can only thrive in an atmosphere that prioritizes, supports and appreciates its importance. An atmosphere of political or cultural intolerance to research has a stifling effect on research efforts. Research needs to be considered a priority for Africa, and African governments need to develop national and regional initiatives that accelerate and support research and research-based education in Africa, in order to build the necessary human capital needed and increase the capacity of institutions to provide valuable training and research opportunities. With enhanced capacity for research, African countries can better decide their research priorities and enforce their own agendas for research, which could help transform societies and facilitate the development of health and development initiatives.

## Acknowledgement

We would like to acknowledge the contribution of two research assistants at CORE Africa (Mumah Sharon and Claude Ngwayu) during the data collection process of this project. We would also like to appreciate the collaboration, advice and support of the CORE Africa consultants.

